# Off-target inhibition by active site-targeting SHP2 inhibitors

**DOI:** 10.1101/295170

**Authors:** Ryouhei Tsutsumi, Hao Ran, Benjamin G. Neel

## Abstract

Due to the involvement of SHP2 (SH2 domain-containing protein tyrosine phosphatase) in human disease, including Noonan syndrome and cancer, several inhibitors targeting SHP2 have been developed. Here, we report that the commonly used SHP2 inhibitor NSC-78788 does not exhibit robust inhibitory effects on growth factor-dependent MAPK (mitogen-activated protein kinase) pathway activation, and that the recently developed active site-targeting SHP2 inhibitors IIB-08, 11a-1, and GS-493 show off-target effects on ligand-evoked activation/trans-phosphorylation of the PDGFRβ (platelet-derived growth factor receptor β). GS-493 also inhibits purified human PDGFRβ and SRC *in vitro*, whereas PDGFRβ inhibition by IIB-08 and 11a-1 occurs only in the cellular context. Our results argue for extreme caution in inferring specific functions for SHP2 based on studies using these inhibitors.

## Introduction

SHP2, encoded by *PTPN11*, is a classic protein-tyrosine phosphatase (PTP), comprising two tandem SH2 domains (N-SH2 and C-SH2, respectively), followed by a catalytic (PTP) domain and a C-terminal tail [1]. In the absence of appropriate cellular stimuli, SHP2 resides in an auto-inhibitory “closed” structure, as a consequence of intra-molecular interactions between the N-SH2 and PTP domains [2]. Upon stimulation of cells with appropriate growth factors, such as platelet-derived growth factor (PDGF) or epidermal growth factor (EGF), the cognate receptor tyrosine kinase (RTK) is activated, resulting in receptor trans-phosphorylation [3, 4], as well as phosphorylation of scaffolding proteins, such as GAB family members [5]. SHP2 binds to phosphorylated RTKs and/or scaffold proteins via SH2 domain/phosphotyrosyl (pY) interactions, which are incompatible with intra-molecular N-SH2/PTP interaction and result in an enzymatically activated “open” structure [1, 2].

Once activated, SHP2 functions as a key positive regulator of RTK-evoked signal transduction, acting upstream of RAS in the ERK MAP kinase pathway [6, 7]. Phosphatase activity is required for SHP2 action [8], although the key substrate(s) remain controversial. In addition, in response to most agonists, SHP2 itself undergoes tyrosyl phosphorylation on two C-terminal tyrosines, which then function as GRB2 binding sites. This “adapter” function for SHP2 enhances RAS/ERK signaling [9].

Mutations that affect residues at the N-SH2/PTP interface of SHP2 cause aberrant enzymatic activation, due to inability to form the “closed” structure. Such mutations are found frequently in, and are responsible for, several disorders, including Noonan syndrome [10], juvenile myelomonocytic leukemia (JMML) [1, 11, 12], and, less frequently, solid tumors such as neuroblastoma [13]. In addition, SHP2, along with its binding partner GAB2, is required for BCR/ABL-evoked transformation and leukemogenesis [1, 14], as well as some forms of breast [1, 15] and other cancers [1]. Recent studies indicate that SHP2 deficiency blocks adaptive resistance to RAF inhibitors [16], and SHP2 is implicated as the major mediator of the programmed cell death 1 (PD-1) [17, 18] and B- and T-lymphocyte attenuator (BTLA) immune checkpoint pathways [19]. For these reasons, SHP2 has emerged as an important drug target, and inhibitors targeting the active site, such as NSC-87877 [20], IIB-08 [21], 11a-1 [22], and GS-493 [23], and more recently, an allosteric inhibitor, SHP099 [24], have been reported.

PTP family members share a conserved PTP catalytic domain [25], so development of active site SHP2 inhibitors has focused largely on achieving specificity versus other PTPs. The above active site-targeting inhibitors, except NSC-87877, reportedly are highly specific for SHP2, compared with other PTPs, including SHP1, which has the highest similarity to SHP2 [21, 22, 23]. However, potential off-target effects of these molecules against other enzyme families (e.g., kinases) have, in general, not been reported.

We tested the effects of the commonly used active site SHP2 inhibitors, NSC-87877, IIB-08, 11a-1, and GS-493 on PDGF signaling in fibroblasts, and found that they either failed to effectively inhibit SHP2 function in cells (NSC-87877) or had off-target effects on protein-tyrosine kinases (PTKs). These results argue strongly that such agents cannot be used alone to infer SHP2 functions in health or disease.

## Materials and Methods

### Cell Culture

*Ptpn11^fl/fl^* MEFs expressing CRE-ER^Tam^ [7], Swiss 3T3 (3T3 Swiss albino) fibroblasts, and HEK293T cells were cultured in Dulbecco-modified Eagle’s medium (DMEM), supplemented with 10% fetal bovine serum (FBS). To induce deletion of *Ptpn11*, *Ptpn11^fl/fl^* MEFs were treated with 1 μM 4-hydroxytamoxifen (4-OHT) for 4 days. All cells tested negative for mycoplasma with the Mycoplasma Plus detection kit (Agilent).

### Antibodies

Mouse monoclonal anti-SHP2 (B-1, sc-7384), anti-PDGFRβ-pY716 (F-10, sc-365464), and anti-ERK2 (D-2, sc-1647), as well as rabbit polyclonal anti-PDGFRβ (958, sc-432) and anti-PDGFRβpY579 (sc-135671) antibodies were purchased from Santa Cruz Biotechnology. Mouse monoclonal anti-MEK1 (61B12, #2352), rabbit monoclonal anti-PDGFRβpY857 (C43E9, #3170), and anti-PDGFRβpY1009 (42F9, #3124), as well as rabbit polyclonal anti-pMEK S217/221 (#9121), anti-pERK1/2 T202/Y204 (#9101), and anti-EGFR (#2232) antibodies were purchased from Cell Signaling. Mouse monoclonal anti-phosphotyrosine antibody cocktail (4G10 platinum, 05-1050) was purchased from Millipore. All antibodies were used at the concentrations recommended by their manufacturers.

### Growth Factors and Chemical Compounds

Recombinant human PDGF-BB and EGF were purchased from Peprotech. 4-OHT was purchased from Sigma. IIB-08 and 11a-1 were provided by Dr. Z.Y. Zhang (Purdue University). GS-493 was provided by Dr. J. Rademann (Freie Universität Berlin) or was purchased from Bioduro. NSC-87877 was purchased from Millipore. SHP099 was purchased from Alputon Inc.

### Expression Constructs, Infection and Sorting

Retroviral expression vectors for wild type SHP2 (WT SHP2) and the mutant SHP2^C459E^ were generated by sub-cloning human *PTPN11* cDNA into pMSCV-IRES-EGFP (Clontech). *Ptpn11^fl/fl^* MEFs expressing WT or mutant SHP2 were generated by retroviral infection, according to the manufacturer’s protocol (pMSCV retrovirus system, Clontech,) followed by fluorescence activated cell sorting (FACS) for EGFP-positive cells.

### Immunoblotting

*Ptpn11^fl/fl^* MEFs expressing CRE-ER^Tam^, Swiss 3T3 fibroblasts, or HEK293T cells were seeded on 6-well plates (MEFs and 3T3 cells, 2 x 10^5^/well; HEK293T cells, 5 x 10^5^/well), followed by serum-starvation (MEFs and 3T3 cells, no FBS; HEK293T cells, no FBS or 0.1% FBS) for 16 h. Starved cells were then stimulated with PDGF-BB (50 ng/ml, final concentration) or EGF (1 ng/ml or 50 ng/ml, final concentration) as indicated. For inhibitor treatment, cells were pre-incubated for 3 h (NSC-87877) or 30 min (other SHP2 inhibitors) in media (1 ml/well) containing the indicated concentrations of each compound with 0.5% DMSO or 0.5% DMSO alone, followed by addition of 10 μl of medium containing 5 μg/ml PDGF-BB (50 ng/ml, final concentration). Cells were lysed in SDS lysis buffer (50 mM Tris-HCl pH7.5, 100 mM NaCl, 1 mM EDTA, 1% SDS, 2 mM Na_3_VO_4_). Lysates were resolved by SDS-PAGE, followed by transfer to Immobilon-FL PVDF membranes (Millipore). Membranes were blocked in 1% BSA/TBS containing 0.1% Tween20 for 30 min, and treated with primary antibodies in blocking buffer for 1 h, followed by treatment with IRDye-conjugated secondary antibodies (LI-COR). Images were obtained by using an ODYSSEY CLx quantitative IR fluorescent detection system (LI-COR) with Image Studio software Ver. 5.2 (LI-COR).

### *In Vitro* Kinase Assays

*In vitro* kinase assays were performed by the SelectScreen™ Biochemical Kinase Profiling Service with the Z’-LYTE Kinase Assay (Thermo). Purified recombinant human PDGFRβ or human SRC were incubated with Z’-LYTE kinase substrate in the presence of serial dilutions of each inhibitor in kinase reaction buffer (50 mM HEPES pH 7.5, 0.01% BRIJ-35, 10 mM MgCl_2_, 2 mM MnCl_2_, 1 mM EGTA, 1 mM DTT) with 100 μM ATP (PDGFRβ) or 50 μM ATP (SRC) at room temperature for 1 h. Relative inhibition for each condition (average of technical duplicates) was calculated by setting the activity in the absence of inhibitor as 0%. IC_50_s were obtained by fitting the data to sigmoidal dose-response curves.

## Results and Discussion

### SHP2 catalytic activity is necessary for PDGF-dependent MAPK activation but not for PDGFR phosphorylation

Exposure of PDGF-BB to cells causes dimerization and activation of PDGFRβ on the plasma membrane, resulting in trans-phosphorylation of multiple receptor tyrosine residues, including Y579, Y716, Y857 and Y1009 [4]. These pYs serve binding sites for SH2-domain or PTB-domain-containing signaling proteins, such as SHP2, which initiate downstream signal transduction [4]. To evaluate the role of SHP2 catalytic activity in PDGF signaling, we employed immortalized mouse embryo fibroblasts (MEFs) from *Ptpn11^fl/fl^* animals, which express Cre-ER^Tam^ [7]. MEFs were reconstituted with wild type (WT) SHP2 or a catalytically inactive SHP2 mutant (C459E), and then were treated with 4-OHT (1 μM) for 4 days to delete endogenous *Ptpn11*. Upon PDGF-BB-treatment (50 ng/ml) after serum-starvation, parental (undeleted) cells showed agonist-evoked phosphorylation/activation of the ERK MAPK pathway components, MEK1 and ERK1/2, whereas these were strongly suppressed by SHP2 depletion (Fig. 1). Suppressed ERK signaling in *Ptpn11* knockout cells was rescued by re-expression of WT SHP2, but not by SHP2^C459E^ (Fig. 1), confirming a requirement for SHP2 catalytic activity in PDGFR-induced ERK activation in fibroblasts. Notably, overall PDGFR tyrosyl phosphorylation, as well as phosphorylation of multiple specific PDGFRβ tyrosyl residues (pY579, pY716, pY857 and pY1009 of PDGFRβ were unaffected, or slightly increased, by the absence of catalytically active SHP2 in *Ptpn11*-knockout cells (Fig. 1). These data indicate that SHP2 does not enhance (and might inhibit some) PDGFRβ tyrosyl phosphorylation events (at these time points) in this immortalized MEF line.

**Fig. 1.**
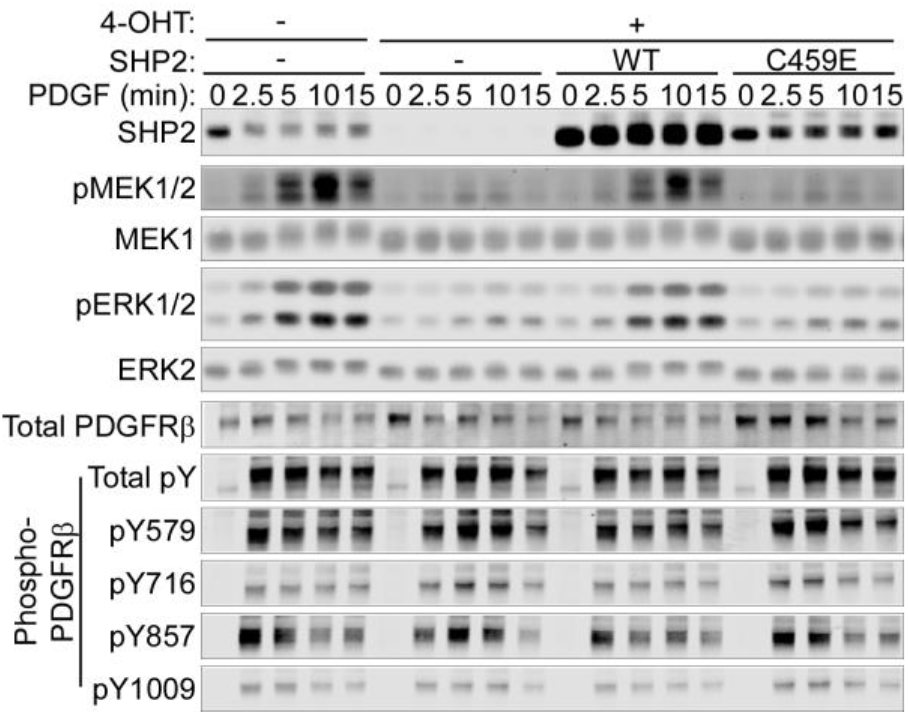
SHP2 catalytic activity is required for PDGF-evoked ERK MAPK signaling, but not for PDGFRβ phosphorylation. *PTPN11^fl/fl^* MEFs expressing CRE-ER^Tam^ with or without retroviral expression of WT or C459E mutant SHP2 were treated with 4-OHT (1 μM) for 4 days or left untreated. Cells were then serum-starved, followed by treatment with PDGF-BB (50 ng/ml) for the indicated times. Lysates were subjected to immunoblotting with the indicated antibodies. Representative immunoblots are shown from one of 3 experiments.

### Active site-targeting SHP2 inhibitors suppress PDGF-evoked ERK activation and PDGFRβ-phosphorylation

We then tested the effects of the active site-targeting SHP2 inhibitors, NSC-87877, IIB-08, 11a-1, and GS-493, at first using Swiss 3T3 fibroblasts. After serum-starvation, cells were pre-treated with the reported inhibitory concentrations of NSC-87877 (20 or 50 μM, 3 h) [20], IIB-08 (10 μM, 30 min) [21], 11a-1 (5 μM, 30 min) [22], or GS-493 (5 μM, 30 min) [23], and then stimulated with PDGF-BB (50 ng/ml, final concentration). NSC-87877 had no detectable effect on ERK MAPK signaling (Fig. 2A). We also failed to reproduce reported inhibitory effects of this compound on EGF (1ng/ml or 50 ng/ml)-stimulated ERK activation in HEK293T cells [20] (Fig. 2B). By contrast, the allosteric SHP2 inhibitor SHP099 effectively suppressed PDGF- or EGF-stimulated ERK MAPK pathway in Swiss 3T3 cells or HEK293T cells, respectively (Fig. 2C).

**Fig. 2.**
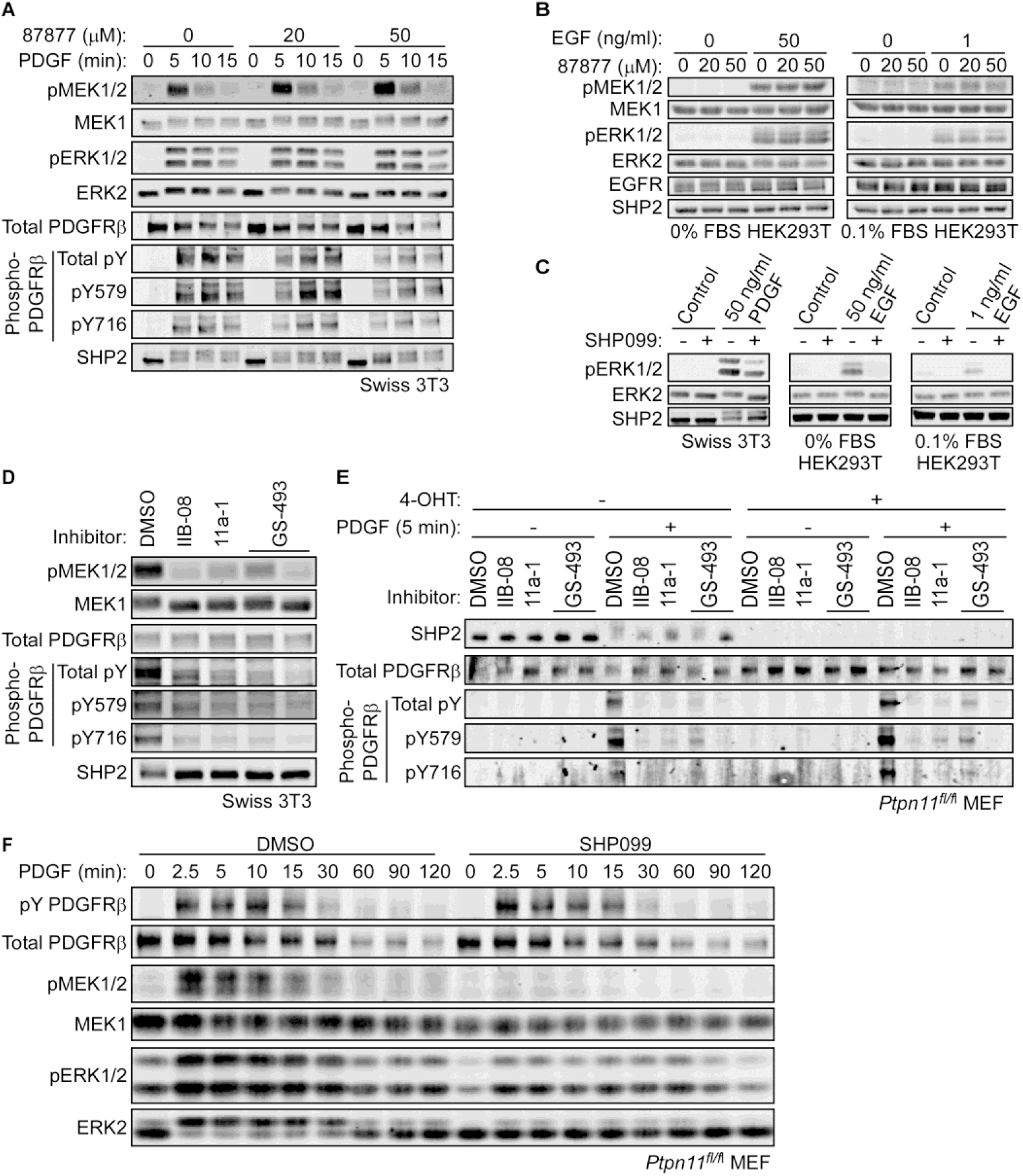
SHP2-independent suppression of PDGFRβ phosphorylation by active site-targeting SHP2 inhibitors. (A) Serum-starved Swiss 3T3 cells were pre-treated with NSC-87877 (20 or 50 μM) for 3 h, followed by addition of PDGF-BB (50 ng/ml, final concentration) for 5 min. (B) HEK293T cells were serum-starved without FBS (left) or with 0.1% FBS (right, according to ref. 20) for 16 h, pre-treated with NSC-87877 (20 or 50 μM) for 3 h, and followed by addition of EGF (50 ng/ml or 1 ng/ml, final concentration, respectively) for 5 min. (C) Serum-starved Swiss 3T3 cells or HEK293T cells were pre-treated with SHP099 (10 μM) for 30 min, followed by addition of PDGF-BB (50 ng/ml) or EGF (50 ng/ml or 1 ng/ml) for 5 min as indicated. Serum-starved Swiss 3T3 cells (D) or *PTPN11^fl/fl^* MEFs expressing CRE-ER^Tam^ treated with or without 4-OHT (E) were pre-treated with IIB-08 (10 μM), 11a-1 (5 μM), GS-493 (5 μM), or DMSO for 30 min, followed by addition of PDGF-BB (50 ng/ml, final concentration) for 5 min. Lysates were then subjected to immunoblotting with the indicated antibodies. Representative immunoblots are shown from one of 2 experiments, respectively. (F) *PTPN11^fl/fl^* MEFs without *PTPN11* deletion were serum-starved and pre-treated with SHP099 (10 μM) for 30 min. Cells were then stimulated with PDGF-BB (50 ng/ml) for the indicated times. Lysates were subjected to immunoblotting with the indicated antibodies. Representative immunoblots are shown from one of 3 experiments.

**Fig. 3.**
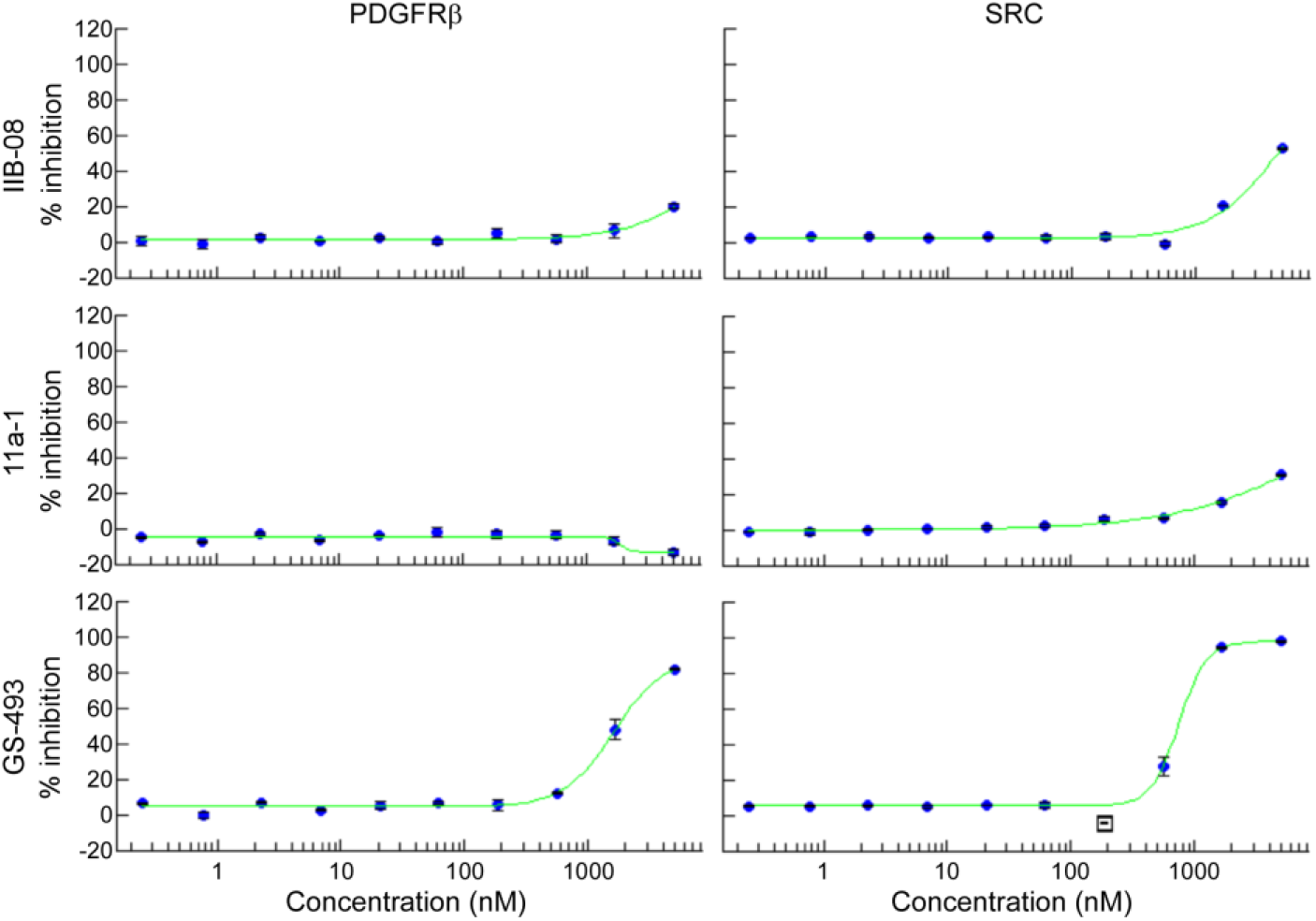
Inhibition of PDGFRβ and SRC *in vitro* by GS-493, but not IIB-08 and 11a-1. Kinase activities of purified human PDGFRβ or human SRC *in vitro* were measured in the presence of serial dilutions of IIB-08, 11a-1, or GS-493. Graphs show relative inhibition of kinase activity in the presence of indicated concentration of each inhibitor (averages of duplicates, blue dots), setting the activity without SHP2 inhibitor as 0%. Data were fit to sigmoidal dose-response curves (green lines). IC_50_s for GS-493 are found in the main text.

The other active site inhibitors tested efficiently abolished PDGF-evoked MEK1 phosphorylation, but also strongly suppressed ligand-dependent tyrosine phosphorylation of PDGFRβ (Fig. 2D). As *Ptpn11* knockout (Fig. 1) did not attenuate PDGFR phosphorylation, we tested these inhibitors in MEFs. In contrast to the effects of SHP2 depletion or replacement of SHP2 with a catalytically inactive SHP2 mutant, IIB-08, 11a-1, and GS-493 suppressed ligand-dependent PDGFRβ phosphorylation in wild type MEFs. Even more importantly, these inhibitors also blocked PDGFRβ phosphorylation in cells lacking SHP2 (Fig. 2E). By contrast, treatment of MEFs with SHP099 suppressed PDGF-dependent MEK/ERK activation without suppressing PDGFRβ phosphorylation (Fig. 2F). These observations show unambiguously that suppression of PDGFRβ phosphorylation by these active site inhibitors is independent of SHP2, and indicate that they have off-target effects, at least in fibroblasts.

### GS-493, but not IIB-08 and 11a-1, inhibits PDGFRβ and SRC kinase activities*in vitro*

To investigate how IIB-08, 11a-1, and GS-493 block PDGFRβ tyrosine phosphorylation, we first asked whether they directly inhibit PDGFRβ kinase activity *in vitro*. Indeed, GS-493 inhibited the purified PDGFRβ kinase domain (IC_50_=1.6 μM), and also inhibited recombinant SRC (IC_50_=746 nM). By contrast, IIB-08 or 11a-1 did not significantly inhibit either isolated kinase domain *in vitro*.

Taken together, our data indicate that NSC-87877—or at least versions of this compound available commercially--lacks clear inhibitory effects on the ERK MAPK pathway, at least in the context of PDGF- or EGF-stimulation of fibroblasts and HEK293T cells. By contrast, GS-493, IIB-08 and 11a-1 suppress PDGFRβ phosphorylation independently of SHP2, and thereby block downstream signaling in PDGF-stimulated cells. GS-493 suppresses ligand-evoked trans-phosphorylation of PDGFRβ likely by directly inhibiting the PDGFRβ kinase domain. GS-493 also inhibits SRC *in vitro*, and given the conservation of kinase domains, might well inhibit other PTKs as well. How IIB-08 and 11a-1 impair PDGFRβ activation in cells, while not inhibiting the PDGFRβ kinase domain *in vitro*, remains unclear. Potential mechanisms include: 1) modulation of receptor affinity for PDGF, 2) inhibition of receptor dimerization or allosteric inhibition via another region of PDGFRβ not included in the recombinant kinase domain; 3) activation of a PTP(s) that dephosphorylates PDGFRβ, or 4) PDGFRβ inhibition via a metabolite(s) generated within cells. As with GS-493, additional potential off-target effects of IIB-08 and 11a-1 on other PTKs cannot, and should not, be excluded.

Inhibiting either SHP2 or the PDGFR can inhibit RAS/ERK/MAPK signaling. Hence, the biological effects of the above inhibitors are difficult, if not impossible, to attribute to SHP2 inhibition. Nevertheless, these inhibitors have been used extensively to probe SHP2 action (reviewed in refs. 26, 27), often as major perturbants of SHP2 function. By contrast, the allosteric inhibitor SHP099 does not have off-target effects on SRC or other tyrosine kinases *in vitro* [24] or in the cellular contexts described here. Even so, comparing the effects of SHP099 and SHP2 depletion (via chemical degradation) indicates that SHP099 also can have off-target effects in at least some settings [28].

In summary, the above study and our results indicate that reports using SHP2 inhibitors as the major means of inferring SHP2 function must be re-evaluated and potentially re-interpreted. For SHP099-like compounds, defined drug-resistant mutants can be used in rescue experiments to demonstrate specificity [24]. Current allosteric SHP2 inhibitors act as “molecular glue” stabilizing the closed, inactive form of the enzyme, and thereby act like “chemical nulls” that potentially block multiple, if not all, aspects of SHP2 function [29]. For this reason, it would certainly be useful to have validated “on-target” catalytic inhibitors. The off-target effects revealed here argue that additional efforts are necessary to achieve this goal, and emphasize the need for careful control experiments and multiple lines of evidence to confidently assign functions to SHP2 using inhibitor approaches.

## Acknowledgements

We thank Drs. Z.Y. Zhang (Purdue University) and J. Rademann (Freie Universität Berlin) for materials. We also thank Ms. X. Wang (Neel lab) for synthesizing plasmids. This work was supported by NIH R37CA49132 to BGN.

## Author contributions

RT and BGN conceived and designed the experiments. RT and HR performed the experiments. RT and BGN wrote the manuscript. BGN supervised the research.

## Abbreviations

4-OHT: 4-hydroxytamoxifen
EGF: epidermal growth factor
MAPK: mitogen-activated protein kinase
PDGF: platelet-derived growth factor
PDGFRβ: platelet-derived growth factor receptor β
PTP: protein-tyrosine phosphatase
PTK: protein-tyrosine kinase
pY: phosphotyrosyl
RTK: receptor tyrosine kinase
SHP2: SH2 domain-containing protein-tyrosine phosphatase

